# Migration of dI5 Reelin-Lmx1b-Zfhx3 and Disabled-1-Lmx1b-Zfhx3 neurons contribute to the superficial dorsal horn and lamina V

**DOI:** 10.64898/2026.03.13.707781

**Authors:** Griselda M. Yvone, Carmine L. Chavez-Martinez, Mahlet A. Mekonnen, Samantha Zimmer, Patricia E. Phelps

## Abstract

In adult superficial dorsal horn, 90% of Reelin (Reln+) and 70% of Disabled-1 (Dab1+) neurons co-express the transcription factor LIM-homeobox 1-beta (Lmx1b+) and therefore are glutamatergic neurons. Here we asked if embryonic Reln+Lmx1b+ and Dab1+Lmx1b+ dorsal horn neurons are derived from Lmx1b-expressing early-born dI5 or late-born dIL_B_ dorsal neurons. On Embryonic day (E)11.5, Reln+ and Dab1+ neurons appear to be part of the migration of early-born dI5 Lmx1b-expressing neurons. Between E12.5-E15.5, the lateral Reln+Lmx1b+ and Dab1+Lmx1b+ neurons migrate circumferentially along the rim of what will become the superficial dorsal horn, whereas medial Reln+Lmx1b+ and Dab1+Lmx1b+ neurons move into the dorsal midline and then migrate into lamina V. The small, late-born dIL_B_ Reln+Lmx1b+ and Dab1-Lmx1b+ neurons fill the superficial dorsal horn. In *Reln* mutants, large Dab1+Lmx1b+ neurons were mispositioned in lamina I and at the border between the superficial and deep dorsal horn. To confirm the identity of the circumferential and midline Reln+Lmx1b+ and Dab1+Lmx1b+ neurons, we asked if they expressed the transcription factor Zfhx3, a marker of dI5 projection neurons. We detected examples of Reln+Lmx1b+Zfhx3+ and Dab1+Lmx1b+Zfhx3+ projection neurons that migrated along the outer rim of the superficial dorsal horn and others that migrated from the midline into lamina V. Taken together, our study demonstrates that the larger Reln+Lmx1b+Zfhx3+ and Dab+Lmx1b+Zfhx3+ neurons represent two subsets of dI5 projections neurons, whereas smaller Reln+Lmx1b+ and Dab1+Lmx1b+ neurons concentrated in lamina II are likely dIL_B_ interneurons.

## Introduction

The *reeler* mouse has a naturally occurring mutation (Falconer, 1951) that encodes a large extracellular matrix-type protein named Reelin (D’Arcangelo et al., 1995). The neuronal positioning errors observed in mice without Reelin (Reln), the two lipoprotein receptors Apolipoprotein E receptor 2 (Apoer2) and Very-low-density-lipoprotein receptor (Vldlr), or Disabled-1 (Dab1) are similar, implicating these proteins in a common signaling pathway (D’Arcangelo et al., 1996; Hiesberger et al., 1999; Howell, Hawkes, et al., 1997; Trommsdorff et al., 1999). After Reln is secreted, it binds to its lipoprotein receptors found on neurons that express the intracellular adaptor protein Dab1. Dab1 is phosphorylated by Src family kinases which activate downstream signaling pathways and induce cytoskeletal changes that influence neuronal migration (Howell, Hawkes, et al., 1997).

The role of Reln signaling in cortical development is widely studied, yet its role in the dorsal spinal cord is still unclear. Initially, we implicated Reln signaling in nociceptive functions as Reln- (Reln+) and Dab1-labeled (Dab1+) neurons are concentrated in dorsal horn areas associated with pain transmission: laminae I-II (superficial dorsal horn), the lateral reticulated area of lamina V, and the lateral spinal nucleus (LSN; Akopians et al., 2008; Villeda et al., 2006). Importantly, adult *Reln^-/-^* and *dab1^-/-^* mice displayed heat hypersensitivity and profound mechanical insensitivity. We also reported that in adult lumbar superficial dorsal horn, about 90% of Reln+ and 70% of Dab1+ neurons co-expressed the transcription factor LIM-homeobox 1-beta (Lmx1b+) and therefore represented separate populations of glutamatergic neurons (Yvone et al., 2020; Yvone et al., 2017). We then detected more Reln+Lmx1b+ and Dab1+Lmx1b+ neurons in adult superficial dorsal horns of *dab1^-/-^* and *Reln^-/-^* compared to their respective wild-type mice. Yet there were fewer Reln+Lmx1b+ and Dab1+Lmx1b+ neurons in *dab1^-/-^* and *Reln^-/-^* lateral lamina V and LSN versus controls (Yvone et al., 2020; Yvone et al., 2017). Ectopic or misplaced neurons within the superficial dorsal horn likely contribute to the nociceptive abnormalities found in *Reln^-/-^* and *dab1^-/-^* mice (Wang et al., 2019; Yvone et al., 2020; Yvone et al., 2017).

During embryonic dorsal spinal cord development, six populations of dorsal interneurons (dI1-6) arise from their respective dorsal progenitor domains. These dorsal interneuron subtypes are defined by transcription factor expression, and migrate to and settle within different spinal cord regions (Gross et al., 2002; Helms & Johnson, 2003; Hernandez-Miranda et al., 2017; Müller et al., 2002). The dI5 neurons migrate laterally, whereas the dIL_B_ neurons migrate into the core of the dorsal horn and differentiate into association interneurons that receive sensory information (Gross et al., 2002; Müller et al., 2002). Notably, the transcription factor Lmx1b marks both the early-born (E10-E12.5) dI5 and late-born (E11-E13) dIL_B_ neurons which are both glutamatergic and required for normal development (Ding et al., 2004; Gross et al., 2002; Lai et al., 2016; Müller et al., 2002; Osseward et al., 2021).

As the majority of Reln+ and Dab1+ neurons in the superficial dorsal horn co-express Lmx1b, we asked whether the Reln+Lmx1b+ and Dab1+Lmx1b+ neurons are subsets of the dI5 or dIL_B_ populations. Initially, we investigated their migratory pathways during dorsal horn development. We then asked how the migrations of Reln+Lmx1b+ and Dab1+Lmx1b+ neurons were altered in *dab1^-/-^* and *Reln^-/-^* mice, respectively and quantified these results. We also asked whether or not the Reln+Lmx1b+ and Dab1+Lmx1b+ neurons were discrete populations of Lmx1b+ neurons as we had observed in adults (Yvone et al., 2020). To establish if the Reln+Lmx1b+ and Dab1+Lmx1b+ neurons were dI5 projection neurons, we asked if these neurons also expressed of the transcription factor Zfhx3 (Osseward et al., 2021). Finally, we correlated our findings to the proposed pattern of dorsal horn development based on spinal cord birthdating studies (Altman & Bayer, 1984; Roome et al., 2020; Sagner et al., 2021)

## Materials and Methods

### Animals

All animal experiments were reviewed and approved by the Chancellor’s Animal Research Committee at UCLA and conducted in accordance with the National Institute of Health guidelines for the care and use of laboratory animals.

#### *Reln* mice

Originally *Reln* (B6C3Fe-ala-*Reln^rl^*) mice were obtained from Jackson Laboratory and a breeding colony was established at UCLA. Mice were genotyped according to D’Arcangelo et al., (1996).

#### *dab1* mice

Mice were obtained from Dr. Brian Howell (SUNY Upstate Medical University, Syracuse, NY) and established as a breeding colony at UCLA. These mice have a *lacZ* reporter fused with the first 22 amino acids of the *dab1* locus, enabling the identification of Dab1 expression with β-gal histochemistry. The generation and characterization of the *dab1^lacZ^* mice were described and genotyped as reported (Abadesco et al., 2014; Pramatarova et al., 2008). In mutants, Dab1 protein expression is eliminated and replaced by β-gal reaction product. The *dab1^lacZ/lacZ^* mice have similar pain behavioral and dorsal horn anatomical abnormalities as *dab1^-/-^* mice (Abadesco et al., 2014; Yvone et al., 2020; Yvone et al., 2017), and therefore the *dab1^lacZ^*mice in this study are identified as *dab1^+/+^* and *dab1^-/-^* mice.

### Tissue preparation and immunohistochemistry

Breeding pairs were checked daily at 8:30 am, and the day of plug detection was established as embryonic day 0.5 (E0.5). Pregnant mice were deeply anesthetized and embryos (E11.5–E15.5) were delivered by Cesarean section. Tail samples were collected for genotyping, and the lower torsos were immersed in 2% PLP (2% Paraformaldehyde; 0.075M Lysine-HCL-monohydride; 0.010M Sodium periodate; 0.1M sodium phosphate) overnight at 4°C. After multiple washes with 0.12 M Millonig’s Phosphate Buffer (97 mM NaOH; 138 mM NaH2PO4), the lumbar enlargement was dissected and cryoprotected in 30% sucrose/Millonig’s for 1-2 days before being frozen in M-1 embedding matrix (Thermo-Scientific, Shandon #1310) and stored at -80°C. Embryonic sections from both genders were used for experiments, sectioned 30 µm thick and slide-mounted.

#### Immunohistochemical procedures

For Reln immunolabeling, goat anti-Reln (1:1,000; R&D Systems; #AF3820) was used together with a Tyramide Signal Amplification (TSA) kit. Sections were first incubated in 1% hydrogen peroxide and 0.1% sodium azide in phosphate-buffered saline (PBS; 0.1M phosphate buffer; 0.9% NaCl) and then blocked in 10% normal donkey serum and 0.1% Triton X-100 in PBS. Sections were incubated with Avidin-Biotin blocking solution (Vector laboratories; kit #SP-2001) before overnight incubation with goat anti-Reln at room temperature. Sections were washed in PBS followed by TNT buffer (0.1M Tris-HCl; 0.15M NaCl; 0.05% Tween-20), and then by a 1 hr incubation with biotinylated horse anti-goat IgG (1:1,000; Vector laboratories; #BA-9500) in TSA-specific blocking buffer (TNB; 0.1M Tris-HCl; 0.15M NaCl; 0.5% Blocking reagent; PerkinElmer #FP1020). Slides were washed with TNT followed by a 1 h incubation with streptavidin-conjugated horseradish peroxidase (1:1,000; PerkinElmer; #NEL750001EA) in TNB, and a 5-min incubation with TSA Plus Fluorescein (1:200; PerkinElmer; #NEL741001KT). Abbreviations for Reln-expressing neurons are single-labeled Reln+, double-labeled Reln+Lmx1b+ or Reln+Zfhx3+, and triple-labeled, Reln+Lmx1b+Zfhx3+ neurons.

For Dab1, Lmx1b, and Zfhx3 labeling, rabbit anti-Dab1 (B3 1:5,000; gift from Dr. Brian Howell, Howell, Gertler, et al., 1997), guinea pig anti-Lmx1b (1:20,000; gift from Drs. Thomas Müller and Carmen Birchmeier, Müller et al., 2002), and sheep anti-Zfhx3 (1:10,000; R&D Systems; #AF7384) were conducted with TSA protocols similar to those reported by (Yvone et al., 2020). Secondary antibodies purchased from Jackson Immunoresearch for TSA experiments included biotinylated donkey anti-guinea pig (#706-065-148), donkey anti-rabbit IgG (#711-065-152), and donkey anti-sheep IgG (#713-065-147). Similar nomenclature is used for single-labeled Dab1+, double-labeled Dab1+Lmx1b+ or Dab1+Zfhx3+, and triple-labeled Dab1+Lmx1b+Zfhx3+ neurons.

All images in the study were obtained with a Zeiss Laser Scanning Microscope (LSM800) with solid-state lasers 488 nm and 640 nm for double-labeled images and 488, 561, and 640 nm for triple-labeled images. For E11.5 and E12.5 images, the entire spinal cord sections were captured with a 20x objective (numerical aperture 0.75), while for E13.5 to E15.5, only dorsal spinal cord sections were imaged. High magnification images were obtained with a 40x oil immersion lens (numerical aperture 1.4) with the pinhole aperture set to 1 Airy unit. Single optical slices are presented for each image in all figures in this study. The images were scanned within the depth of the section where signals of all channels were detected to generate z-series of approximately 4-7 µm, with a z-separation of 1 µm. ZEN (Zeiss Efficient Navigation) lite imaging software was used for obtaining images, which were then transferred to Photoshop for figure assembly.

### Statistical Analyses

To compare the migration patterns of Dab1+Lmx1b+ dorsal spinal cord neurons between genotypes at E13.5-E15.5, we quantified the number of Dab1+Lmx1b+ neurons from 1 µm optical sections of *Reln^+/+^* and *Reln^-/-^* embryos (n=3 embryos per genotype at each age, with 3-6 hemisections analyzed per embryo). We divided each dorsal horn hemisection into 12 equal-sized grids. The dorsoventral border included laminae I-V, while the mediolateral border extended from the ventricular zone midline to the lateral spinal cord edge. Results are presented as an average percent of total Dab1+Lmx1b+ cells recorded for each grid per age. Average percentages were analyzed by a two-way analysis of variance (ANOVA; genotype and grid as factors), followed by a Holm-Sidak *post hoc* test. Statistical analyses were performed with GraphPad Prism 9.

We also quantified the extent to which the Reln+Lmx1b+ and Dab1+Lmx1b+ dorsal midline neurons co-expressed Zfhx3 in E14.5 wild-type embryos (n=3). A box, 100 µm x 100 µm in size, was drawn surrounding the center of the cluster of neurons in the dorsal midline above the ventricular zone at E14.5. In sections processed for Reln, Lmx1b and Zfhx3, we counted Reln+Lmx1b+Zfhx3+, Reln+Zfhx3+, Lmx1b+Zfhx3+, and Zfhx3+ neurons from 1 µm optical sections and calculated the percentage of each cell type. After Dab1, Lmx1b and Zfhx3 localization, Dab1+Lmx1b+Zfhx3+, Dab1+Zfhx3+, Lmx1b+Zfhx3+, and Zfhx3+ neurons were counted and results analyzed as above.

## Results

### Early Reln+ dorsal horn neurons appear to be Lmx1b+ dI5 neurons

As the majority of Reln+ neurons in the adult superficial dorsal horn co-expressed Lmx1b, and these neurons are mispositioned in *dab1^-/-^* mice (Yvone et al., 2020) we asked if Reln+Lmx1b+ neurons were part of the early-born dI5 population in *dab1* mice. At E11.5, Reln expression (green) was concentrated ventrally and laterally, and axons of Reln-expressing commissural neurons crossed in the ventral commissure (VC; Fig. 1A, B). As reported, Lmx1b expression marked the nuclei of the dI5 postmitotic neurons (red; Ding et al., 2004; Gross et al., 2002; Müller et al., 2002) and labeled the floor plate (Fig. 1A1, B1). Lmx1b+ (Fig. 1A1, B1) and Reln+ neurons (Fig. 1A, B) were found together with the Reln+Lmx1b+ cells (Fig. 1A3, B3) in both genotypes. Because the Lmx1b+ dorsal spinal cord neurons present at E11.5 are predominantly in the dI5 group (Gross et al., 2002; Müller et al., 2002), the Reln+Lmx1b+ neurons concentrated dorsolaterally may be dI5 neurons (Fig. 1A2-3, B2-3).

**Figure 1.**
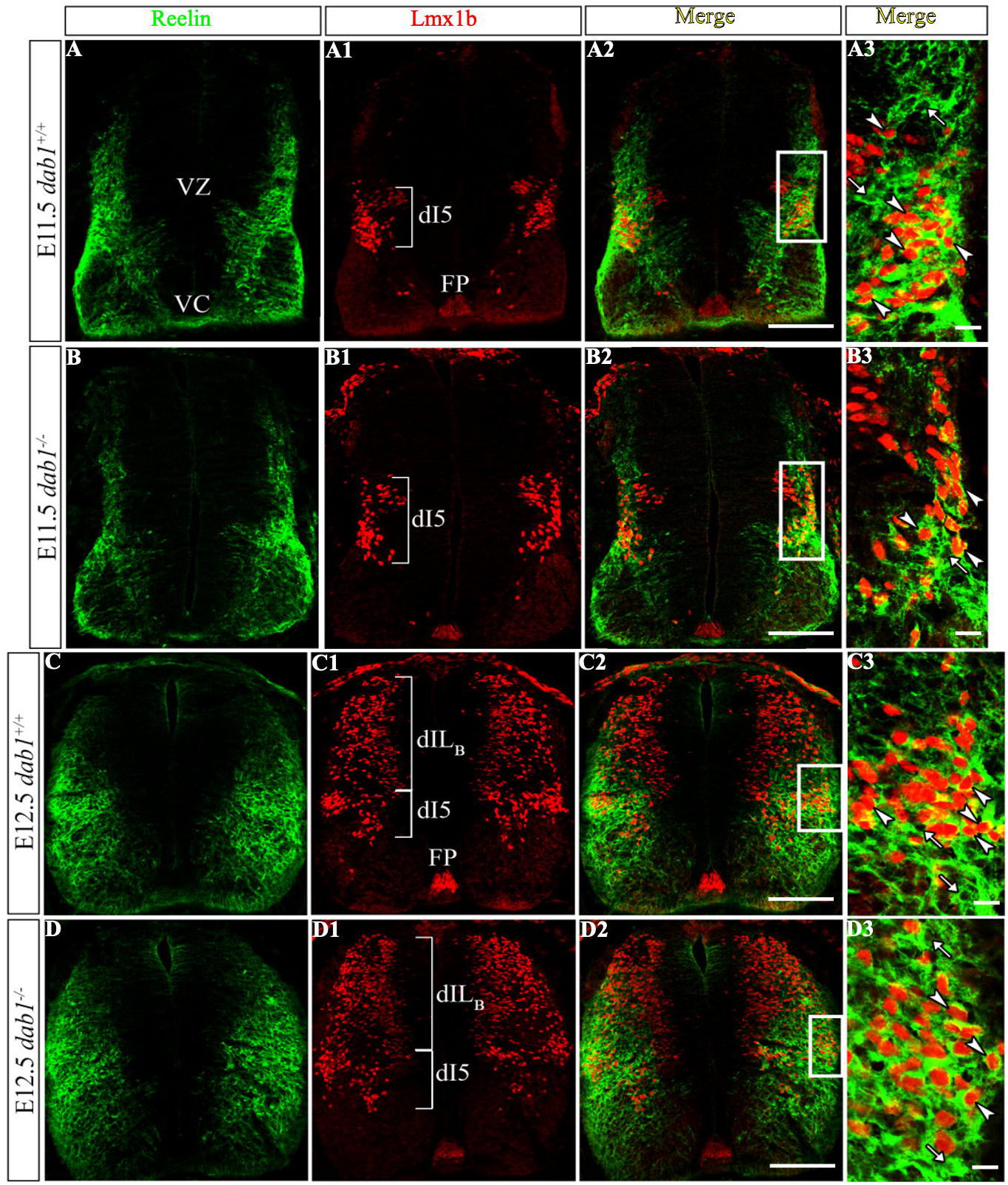
Some dI5 Lmx1b+ neurons co-express Reln in E11.5-E12.5 spinal cord. Single optical sections show Reln+ (green) and Lmx1b+ (red nuclei) co-localization at E11.5 (***A-B***) and E12.5 (***C-D***) in *dab1^+/+^* (***A-A3; C-C3***) and *dab1^-/-^* (***B-B3; D-D3***) spinal cords. Boxes in ***A2-D2*** are enlarged in ***A3-D3***. Reln+Lmx1b+ cells are marked with white arrowheads, and Reln+ cells, with white arrows. ***A-B,*** At E11.5, Reln immunoreactivity (***A, B***) is high along the dorsolateral spinal cord, and in the ventral commissure (VC). The ventricular zone (VZ) is unlabeled. Lmx1b expression (***A1, B1***) is localized laterally in the early-born dI5 group and the floor plate (FP). Reln+Lmx1b+ neurons are within the lateral-most cluster of dI5 neurons (***A2-3; B2-3***). ***C-D***, By E12.5, Reln expression (***C, D***) has expanded dorsally and medially. The lateral Lmx1b+ dI5 neurons are clustered (***C1***, ***D1***) whereas the dIL_B_ group is broadly distributed in the dorsal spinal cord (***C1, D1***). Most Reln+Lmx1b+ neurons are found laterally, as part of the dI5 cohort (***C2-3; D2-3***). Scale bars: ***A-D, A1-D1, A2-D2,*** 100 µm; ***A3-D3,*** 20 µm.

**Figure 2.**
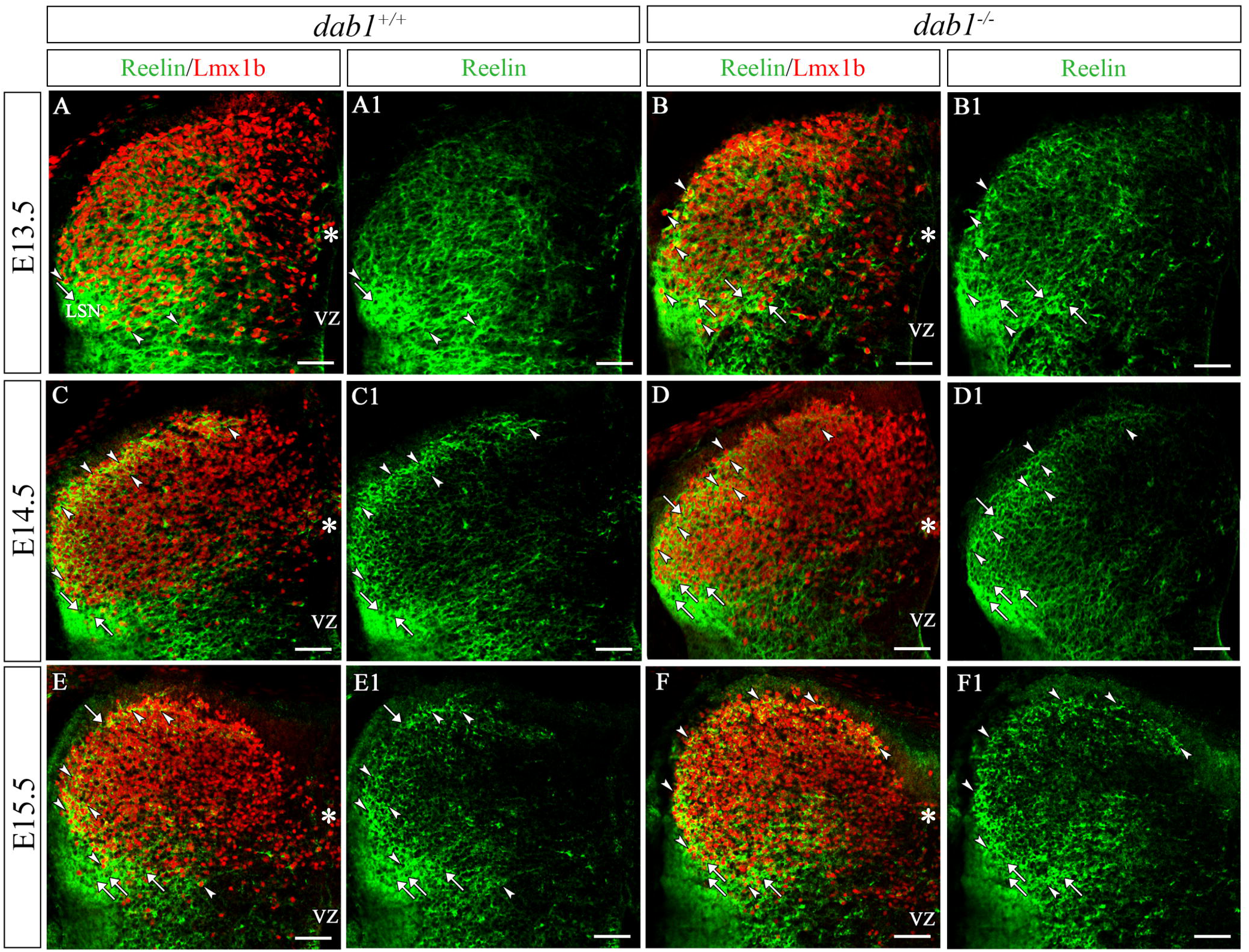
Reln+ neurons appear to migrate with subsets of Lmx1b+ dI5 and dIL_B_ neurons. Reln+ (green) and Lmx1b+ (red nuclei) neurons in E13.5 (***A-B***), E14.5 (***C-D***) and E15.5 (***E-F***) dorsal horns of *dab1^+/+^* (***A-A1; C-C1; E-E1***) and *dab1^-/-^* mice (***B-B1; D-D1; F-F1***). ***A-B***, Numerous Lmx1b+ neurons fill the E13.5 dorsal horn, most of which are part of the late-born dIL_B_ cohort. Reln+ (white arrows) and Reln+Lmx1b+ neurons (white arrowheads) are concentrated laterally and others are found near the midline (white asterisks). Reln is more broadly expressed in the dorsal horn at E13.5 than at E14.5-15.5. Reln+Lmx1b+ cells are detected along the rim of the *dab1^-/-^*dorsal horn (***B, B1***). Reln is highly expressed in the lateral spinal nucleus (LSN). ***C-D,*** At E14.5, Reln+ and Reln+Lmx1b+ neurons are positioned in the rim of the *dab1^+/+^* and *dab1^-/-^* superficial dorsal horn. A few medially located Reln+Lmx1b+ neurons (white asterisks in ***C-F***) are found in both genotypes. ***E-F***, E15.5 Reln+ and Reln+Lmx1b+ neurons are concentrated in the superficial dorsal horn, in the LSN, and in the adjacent reticulated area of lateral lamina V. Scale bars: ***A-F, A1-F1,*** 50 µm.

**Figure 3.**
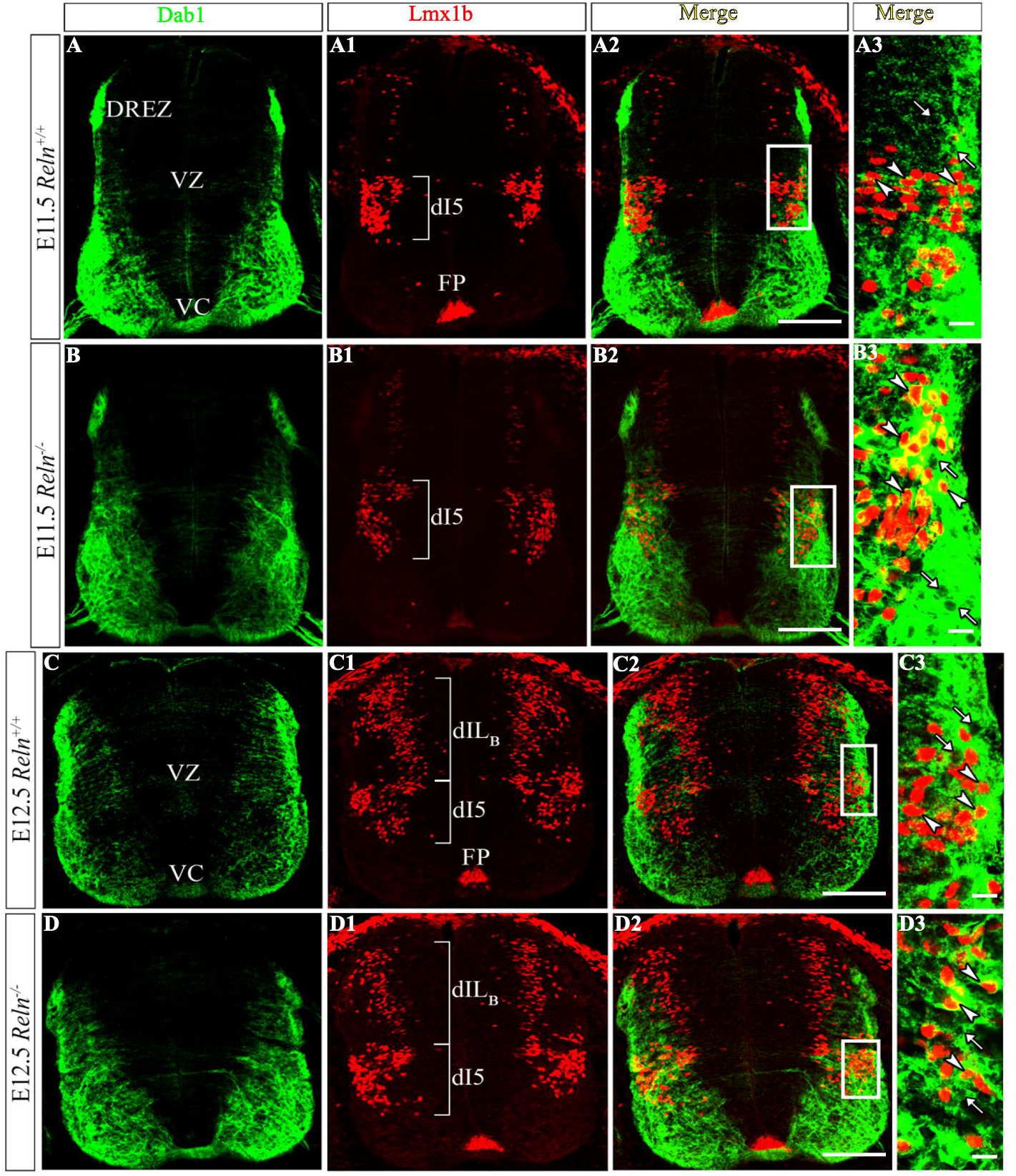
Some Dab1+ dorsal horn neurons are part of the Lmx1b+ dI5 population. Dab1+ (green) and Lmx1b+ (red nuclei) neurons at E11.5 (***A, B***) and E12.5 (***C, D***) of *Reln^+/+^* (***A-A3; C-C3***) and *Reln^-/-^* (***B-B3; D-D3***) spinal cords. Boxes in ***A2-D2*** are enlarged in ***A3-D3***. ***A-B,*** At E11.5, Dab1 immunoreactivity (***A, B***) marks somatic motor neurons, axons in the ventral commissure (VC), and the putative dorsal root entry zone (DREZ). Lmx1b expression (***A1, B1***) is concentrated in dI5 neurons and the floor plate (FP). Dab1+Lmx1b+ neurons (white arrowheads) are in the lateral dI5 group and Dab1+ neurons (white arrows) are present (***A2-3; B2-3***). ***C-D***, At E12.5, the lateral dI5 and broadly dispersed dIL_B_ groups are evident in both genotypes (***C1, D1***). Dab1 and Lmx1b expression patterns appear similar in both genotypes. Scale bars: ***A-D; A1-D1, A2-D2,*** 100 µm; ***A3-D3,*** 20 µm.

At E12.5, Reln expression had expanded further dorsally, ventrally, and medially (Fig. 1C, D; Kubasak et al., 2004). Reln+ and Reln+Lmx1b+ neurons were detected among the lateral dI5 Lmx1b+ neurons (Fig. 1C2-3, D2-3). At both ages, the patterns of Reln+ and Reln+Lmx1b+ neurons appeared similar between the genotypes (Fig. 1A-D, A2-D2). The dIL_B_ population that expresses Lmx1b appeared at E12.5, is derived from a broad progenitor domain, and populated much of the dorsal horn (Fig. 1C1, D1; Gross et al., 2002; Müller et al., 2002). At these early ages, the Reln+Lmx1b+ neurons appeared to associate primarily with the dI5 group.

### Reln+Lmx1b+ neurons contribute to the superficial dorsal horn and the LSN

Compared to both earlier and later ages, Reln expression at E13.5 was broadly detected throughout the dorsal horn (Fig. 2A1, B1). Of the Reln+ and Reln+Lmx1b+ neurons concentrated near the lateral dorsal horn (Fig. 2A-A1, B-B1), some began to migrate circumferentially along the rim of the E13.5 dorsal horn in both genotypes, and they continued their circumferential migration at E14.5-E15.5 (Fig. 2A-F). The highest concentration of Reln expression at these ages, however, remained adjacent to the border of the lateral curve of the nascent dorsal horn, where the LSN and the lateral reticulated area of lamina V are located (Fig. 2C-F). This supports our previous observation that Reln expression in the lateral marginal zone (future lateral funiculus) is primarily secreted Reln (Kubasak et al., 2004). When we blocked Reelin secretion with Brefeldin-A in embryonic organotypic cultures, slices treated with Brefeldin-A had little evidence of secreted Reln in the lateral marginal zone or LSN compared to controls (Kubasak et al., 2004).

An extraordinary expansion of Lmx1b expression characterizes the dIL_B_ population at E13.5 (Fig. 2A, B; Gross et al., 2002; Müller et al., 2002) and Reln expression is broadly distributed in the dorsal horn (Fig. 2A1, B1). By E14.5-15.5 Reelin immunoreactivity is concentrated around Lmx1b-labeled nuclei within the superficial dorsal horn (Fig. 2C-F). In adults, there are many small Reln+Lmx1b+ cells in laminae I-II (Yvone et al., 2020) which likely originated from the dIL_B_ group.

At E13.5-E14.5, we also detected medially-located Reln+Lmx1b+ neurons (Fig. 2A-D, asterisks) which were near or dorsal to the ventricular zone (Fig. 2C-C1, D-D1, asterisks). By E15.5, only a few of the medial Reln+Lmx1b+ neurons remained near the ventricular zone (Fig. 2E-F, asterisks), as most of these neurons appeared to migrate laterally. Based on their initial medial location, we could not distinguish if they were derived from dI5 or dIL_B_ clusters.

### Early Dab1+ dorsal horn neurons appear to be part of the Lmx1b+ dI5 neurons

Because 70% of adult Dab1+ laminae I-II neurons also express Lmx1b, and Dab1+Lmx1b+ neurons are mispositioned in adult *Reln^-/-^* mice (Yvone et al., 2017), we also examined the developmental origins of these neurons. At E11.5 and E12.5, Dab1 expression was detected in somatic motor neurons (Abadesco et al., 2014; Palmesino et al., 2010), in neurons with axons that cross in the ventral commissure (VC), and in some sensory afferents with axons that project into the putative dorsal root entry zone (DREZ; Fig. 3A-B). At both ages, Dab1+ and Dab1+Lmx1b+ neurons were associated with the lateral-most part of the dorsal horn in both genotypes (Fig. 3A2,3 – 3D2,3). The Dab1+Lmx1b+ neurons occupy the lateral part of the E11.5-12.5 dorsal spinal cord (Fig. 3, Gross et al., 2002; Müller et al., 2002), and these neurons appear to contribute to the dI5 group.

### Dab1+Lmx1b+ neurons exhibit migratory errors in *Reln^-/-^* superficial dorsal horn

At E13.5, Dab1+Lmx1b+ neurons were detected along the dorsolateral rim of the superficial dorsal horn, while Lmx1b+ neurons filled the core of both *Reln^+/+^* and *Reln^-/-^* dorsal horns (Fig. 4A-A1, B-B1). Laterally-located Dab1+Lmx1b+ cells in both genotypes were present as well as Dab+ neurons (Fig. 4A, B). The Dab1+ and Dab1+Lmx1b+ neurons detected along the outer rim of the lateral dorsal horn at E13.5 (Fig. 4A-A1) were found further medially at later stages (Fig. 4C-F).

**Figure 4.**
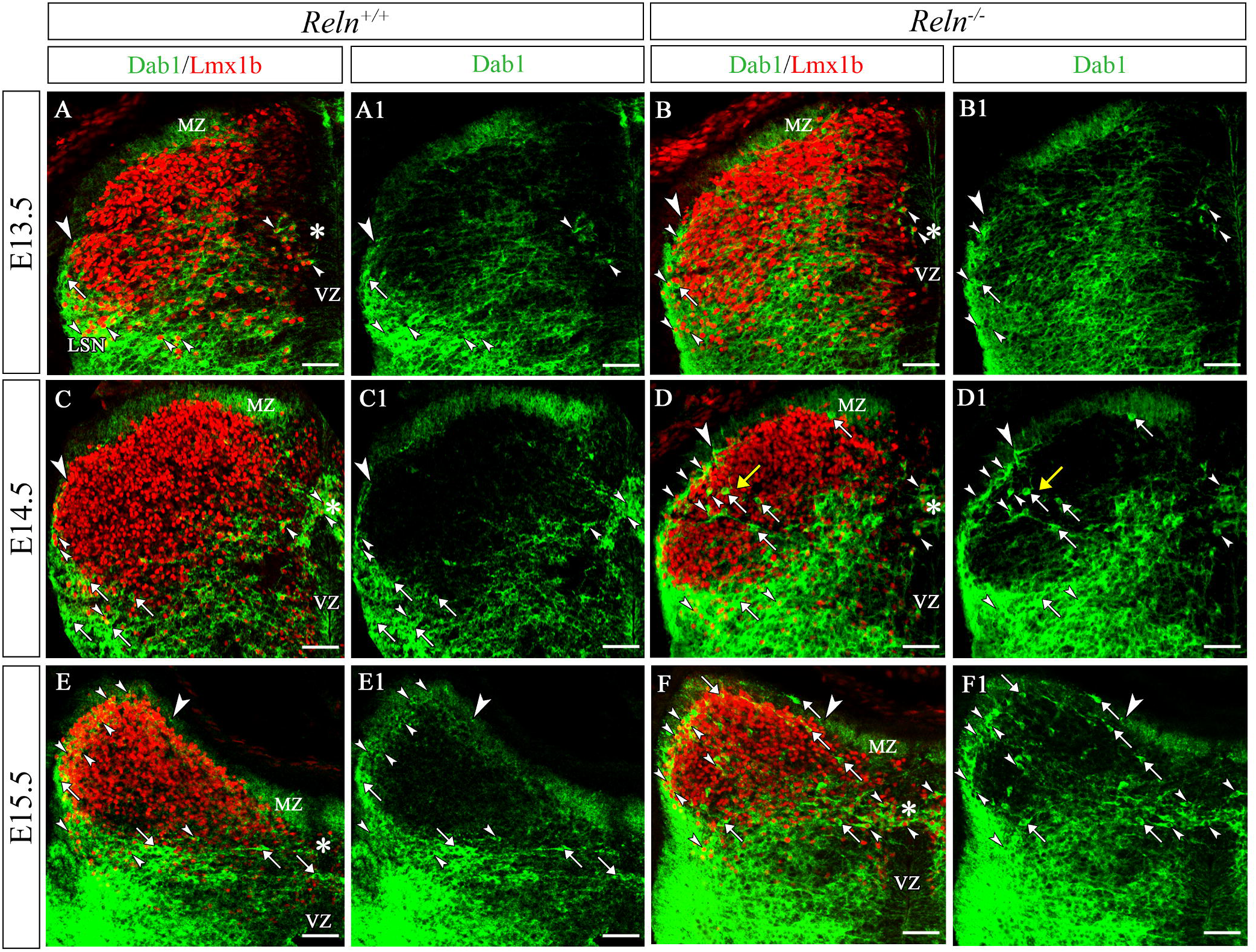
Dab1+ and Dab1+Lmx1b+ neurons display migratory errors in the absence of Reln. Dab1+ (green) and Lmx1b+ (red nuclei) expression patterns in *Reln^+/+^* (***A-A1; C-C1; E-E1***) and *Reln^-/-^* (***B-B1; D-D1; F-F1***) dorsal horns at E13.5 (***A-B***), E14.5 (***C-D***) and E15.5 (***E-F***). The highest concentration of Dab1 is found just lateral to the curve of the nascent superficial dorsal horn, near the lateral spinal nucleus (LSN). Dab1+ axons detected near the dorsal midline are part of the dorsal marginal zone (MZ, ***A-F).*** Large white arrowheads in ***A-F*** mark the furthest extent of the circumferential migrations of Dab1+ and Dab1+Lmx1b+ neurons. ***A-B***, At E13.5, Dab1+Lmx1b+ neurons are found laterally (small arrowheads) together with a few Dab1+ neurons (arrows). Dab1+Lmx1b+ neurons are also found near the midline (asterisks near ventricular zone, VZ). ***C-D,*** At E14.5, Dab1+ (white arrows) and Dab1+Lmx1b+ neurons (white arrowheads) are found laterally and in the midline (asterisks) dorsal to the VZ of *Reln^+/+^* (***C, C1***) and *Reln^-/-^*mice (***D, D1***). Dab1+ and dI5 Dab1+Lmx1b+ neurons are positioned further medially along the rim of the superficial dorsal horn in the *Reln^-/-^* (***D, D1;*** 1 large and 3 small white arrowheads) than in *Reln^+/+^* mice (***C, C1;*** large white arrowhead). Migratory errors are detected in *Reln^-/-^*dorsal horn. The core of the *Reln^+/+^* dorsal horn is almost devoid of Dab1+ neurons, whereas Dab1+ and Dab1+Lmx1b+ neurons coursed tangentially across this area in *Reln^-/-^* mice (***D, D1;*** yellow arrow, enlarged in Supplemental Figure 2). A thickened lamina I also shows some out-of-place dI5 neurons. ***E-F,*** Dab1+ and Dab1+Lmx1b+ neurons in E15.5 *Reln^+/+^* mice are detected within the rim of the superficial dorsal horn and in lamina V (***E, E1***). Dab1+ and Dab1+Lmx1b+ neurons are found within the core of the *Reln^-/-^*, but not in *Reln^+/+^* dorsal horn. Scale bars: ***A-F, A1-F1,*** 50 µm.

To confirm differences in the migratory patterns between *Reln^+/+^* and *Reln^-/-^* embryos, we quantified the number of Dab1+Lmx1b+ dorsal horn neurons from single optical slices by dividing the dorsal spinal cord into 12 equally-sized grids (Fig. 5A). Results show the average percentage of total Dab1+Lmx1b+ cells located in each grid. At E13.5, some of the Dab1+Lmx1b+ neurons are localized in the lateral grids (Fig. 5A, B, grids 5 and 9), and over the next two days, appear to migrate circumferentially and settle along the rim of the superficial dorsal horn (Fig. 5C-F, grids 1, 2, 5). Developmental differences in Dab1+Lmx1b+ neurons between *Reln^+/+^* and *Reln^-/-^* mice were most distinct at E14.5. Dab1+Lmx1b+ neurons had now advanced further dorsomedially along the outer rim of the dorsal horn. Compared to the E14.5 *Reln^+/+^* (Fig. 4C, C1), *Reln^-/-^* mice had a thicker band of Dab1+Lmx1b+ lamina I cells with some cells mispositioned (Fig 4D, D1). Our quantification confirmed a lower percentage of Dab1+Lmx1b+ neurons in *Reln^+/+^* versus *Reln^-/-^*grid 5 (Fig. 5, *Reln^+/+^* 10.2%±4.6% vs. *Reln^-/-^* 20.5%±3%, *p=*0.0116). In addition, there were Dab1-immunoreactive axons, partially derived from the dorsal root entry zone, that accumulated in the marginal zone (MZ) along the medial rim of the dorsal horn in both genotypes and contributed to the dorsal columns (DC; Fig. 4, Supplemental Figure 1).

**Figure 5.**
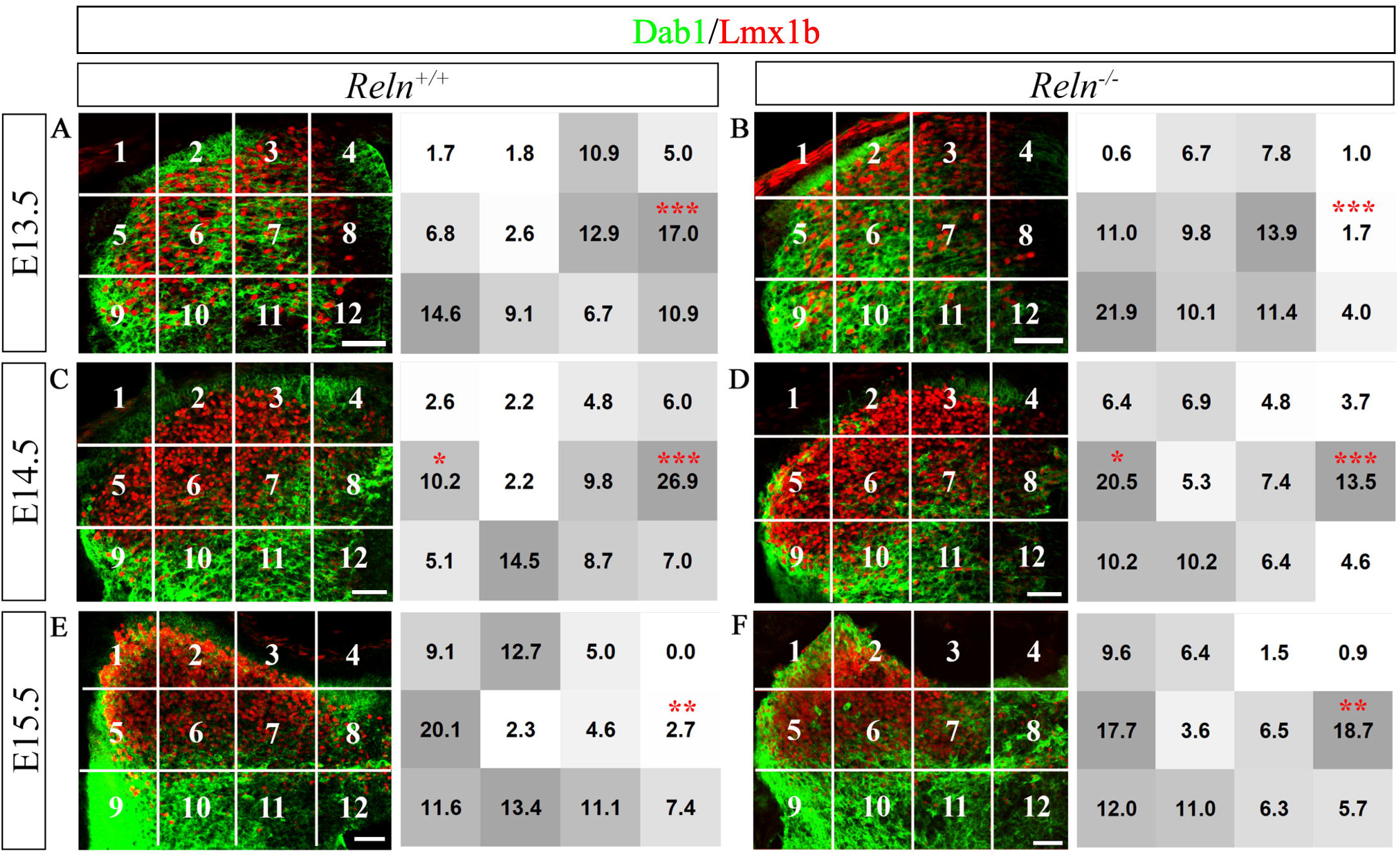
Statistical analyses reveal differences in neuronal positioning of Dab1+Lmx1b+ neurons between genotypes. Each dorsal horn hemisection was subdivided into 12 equal-sized grids. Grids 1, 5, and 9 were assigned to the lateral dorsal horn and grids 4, 8, and 12 to the midline. The percentage of Dab1+Lmx1b+ neurons within each grid was evaluated from single optical slices of *Reln^+/+^* and *Reln^-/-^* dorsal horn and compared between genotypes. ***A, B***, At E13.5, there is a higher percentage of Dab1+Lmx1b+ neurons in grid 8 of *Reln^+/+^* than found in *Reln^-/-^* mice. ***C, D,*** On E14.5, there is a higher percentage of Dab1+Lmx1b+ neurons in the *Reln^+/+^* midline grid 8 than in *Reln^-/-^*mice, and the *Reln^-/-^* mice have more double-labeled neurons in grid 5 than *Reln^+/+^* mice. ***E, F,*** By E15.5, the percentage of Dab1+Lmx1b+ neurons in grid 8 is higher in *Reln^-/-^* than in *Reln^+/+^* mice. Sample: n=3 embryos per genotype per age, with 3 hemisections analyzed per embryo. Significance is represented as *p < 0.05, **p < 0.01, or ***p < 0.001. Scale bars: ***A-F,*** 50 µm.

### Dab1+Lmx1b+ neurons show migratory errors in *Reln^-/-^* deep dorsal horn

A major migratory error was obvious at E14.5 as the *Reln^+/+^* deep dorsal horn was almost completely devoid of Dab1 expression (Fig. 4C1), whereas in the *Reln^-/-^* mice, we consistently detected mispositioned Dab1+ and Dab1+Lmx1b+ neurons (Fig. 4D1, yellow arrow). The misplaced neurons in *Reln^-/-^* mice migrated tangentially across the otherwise Dab1-negative deep dorsal horn (Fig. 4D-D1; Supplemental Figure 2). Examples of tangentially migrating Dab1+ and Dab1+Lmx1b+ neurons also were present in E15.5 *Reln^-/-^* mice (Fig. 4F-F1).

A second prominent group of Dab1+Lmx1b+ neurons was found near the ventricular zone in both genotypes between E13.5-E15.5 (Fig. 4A-F; asterisks). We found a higher percentage of Dab1+Lmx1b+ cells near the ventricular zone in E13.5 *Reln^+/+^* compared to *Reln^-/-^*mice (Fig. 5A-B, grid 8: 17%±3% vs. 1.7%± 0.9%, *p*=0.0001). By E14.5, these large Dab1+Lmx1b+ neurons formed a distinct cluster in the midline just dorsal to the receding ventricular zone (Fig. 6A-A3, B-B3), and a higher percentage of these neurons were found in *Reln^+/+^* compared to *Reln^-/-^* mice (Fig. 5C-D; grid 8: 26.9%±1.3% vs. 13.5%±3.3%, *p*=0.0004). The Dab1+ and Dab1+Lmx1b+ neurons in E14.5 *Reln^-/-^* (Fig. 6B-B3) appeared disorganized in the midline compared to those in *Reln^+/+^* embryos (Fig. 6A-A3). By E15.5, Dab1+Lmx1b+ neurons in *Reln^+/+^* mice formed a stream of cells that migrated into the deep dorsal horn (Figs. 4E-E1; 6C-C2), whereas in *Reln^-/-^* mice, more of these neurons were still found medially (Figs. 4F-F1; 6D-D2). Consistent with these observations, fewer cells remained in the dorsal midline of E15.5 *Reln^+/+^* than in *Reln^-/-^* mice (Fig. 5E vs F, grid 8, 2.7±1% vs. 18.7%± 4%, *p*=0.001). Thus, we found evidence of mispositioning errors in Dab1+Lmx1b+ cells in the dorsal midline of *Reln^-/-^* mice and in their migration into presumptive lamina V. Our results also showed a major migratory error which led to incorrectly positioned Dab1+ and Dab1+Lmx1b+ neurons near the border between the *Reln^-/-^* laminae II and III.

**Figure 6:**
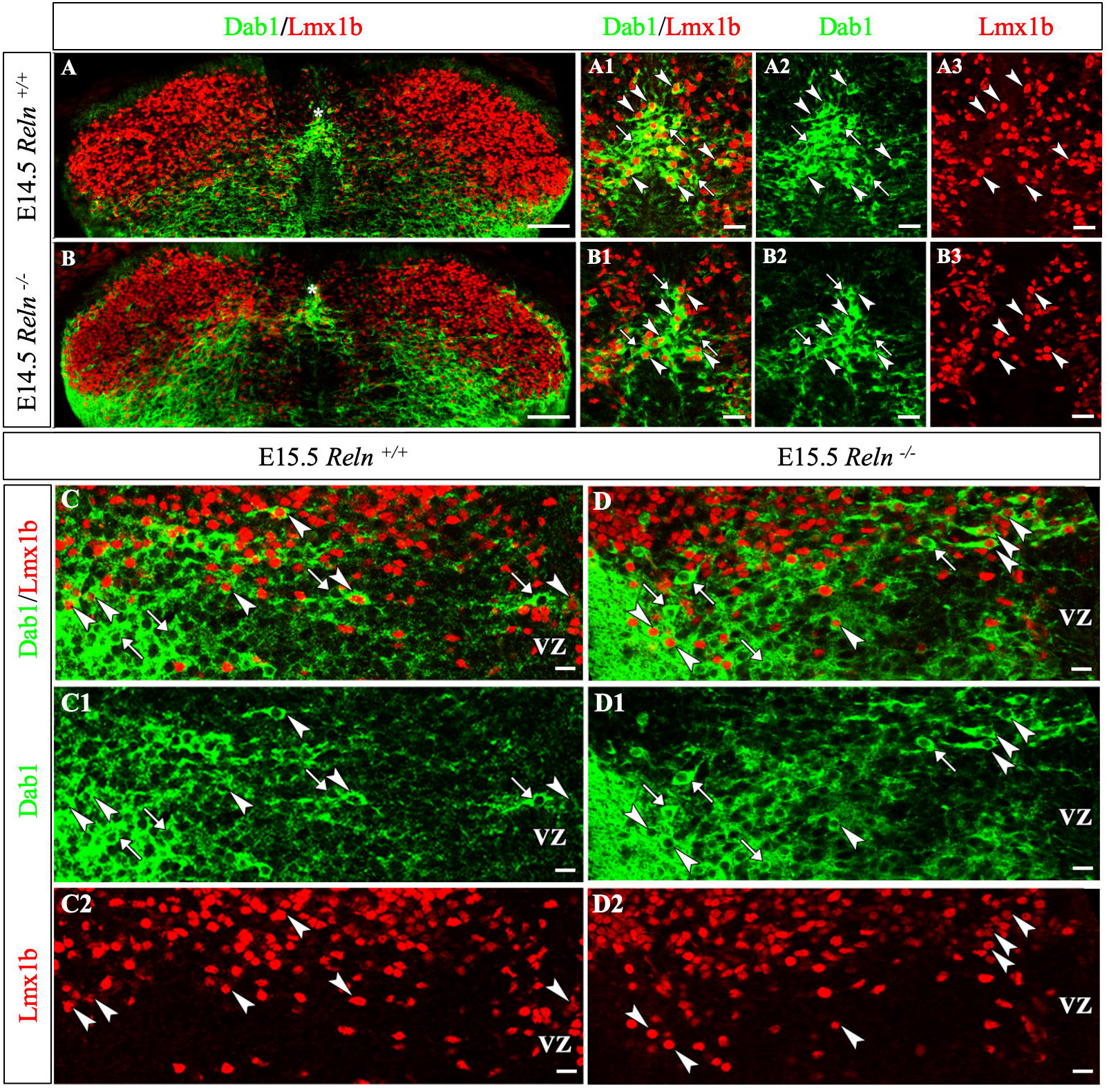
Compared with *Reln^+/+^* mice, Dab1+ and Dab1+Lmx1b+ neurons appear unevenly distributed in the *Reln^-/-^*midline and putative lamina V. Dab1+ (green) and Dab1+Lmx1b (green cytoplasm with red nuclei) expression patterns in *Reln^+/+^* (***A-A3; C-C2***) and *Reln^-/-^* (***B-B3; D-D2***) dorsal horns at E14.5 (***A-B)*** and E15.5 (***C-D***). ***A-B,*** E14.5 Dab1+Lmx1b+ neurons are located in the midline (white asterisks), just dorsal to the receding ventricular zone (VZ). Enlargements of these midline cells (***A1-A3; B1-B3***) show that Dab1+Lmx1b+ neurons (white arrowheads) appear more symmetrically distributed in *Reln^+/+^* (***A-A3***) than in *Reln^-/-^* mice (***B-B3***). A few Dab1+ neurons (white arrows) are present. ***C-C2,*** E15.5 *Reln^+/+^* images enlarged from figure 4E. Dab1+ (white arrows) and Dab1+Lmx1b+ neurons (white arrowheads) migrate in an organized stream into lamina V. ***D-D2,*** Images enlarged from figure 4F. This migration in a *Reln^-/-^* mouse appears disorganized as more Dab1+ and Dab1+Lmx1b+ neurons are clustered medially than in *Reln^+/+^* dorsal horn. Scale bars: ***A, B,*** 50 µm; ***A1-A3, B1-B3,*** 25 µm; ***C-C2, D-D2,*** 20 µm.

### Reln+Lmx1b+ and Dab1+Lmx1b+ neurons in the superficial dorsal horn are independent populations but some Dab1+Lmx1b+ lamina V cells may contain Reelin

Because double *Apoer2/Vldlr* mutant mice express high levels of both Reln and Dab1 (Rice et al., 1998; Trommsdorff et al., 1999), we previously examined 1-month-old double-receptor wild-type (*ApoER2^+/+^/Vldlr^+/+^)* and knock out (*ApoER2^-/-^/Vldlr^-/-^)* mice to determine if the Reln+Lmx1b+ and Dab1+Lmx1b+ neurons were distinct populations in the superficial dorsal horn, and concluded that they were different subsets of Lmx1b neurons (Yvone et al., 2020). Here we performed triple labeling of Dab1, Reln, and Lmx1b at E12.5 and E14.5 on *dab1^+/+^*mice (Fig. 7A-G) to confirm that early embryonic Reln+Lmx1b+ and Dab1+Lmx1b+ neurons are separate subsets of Lmx1b neurons. At E12.5, their locations are partially intermixed, with Reln+Lmx1b+ neurons located more ventrally (Fig. 7A, B, C1, D1) and the Dab1+Lmx1b+ neurons found further dorsally (Fig. 7A, B, C, D). They did not appear to co-localize (Fig. 7B).

**Figure 7:**
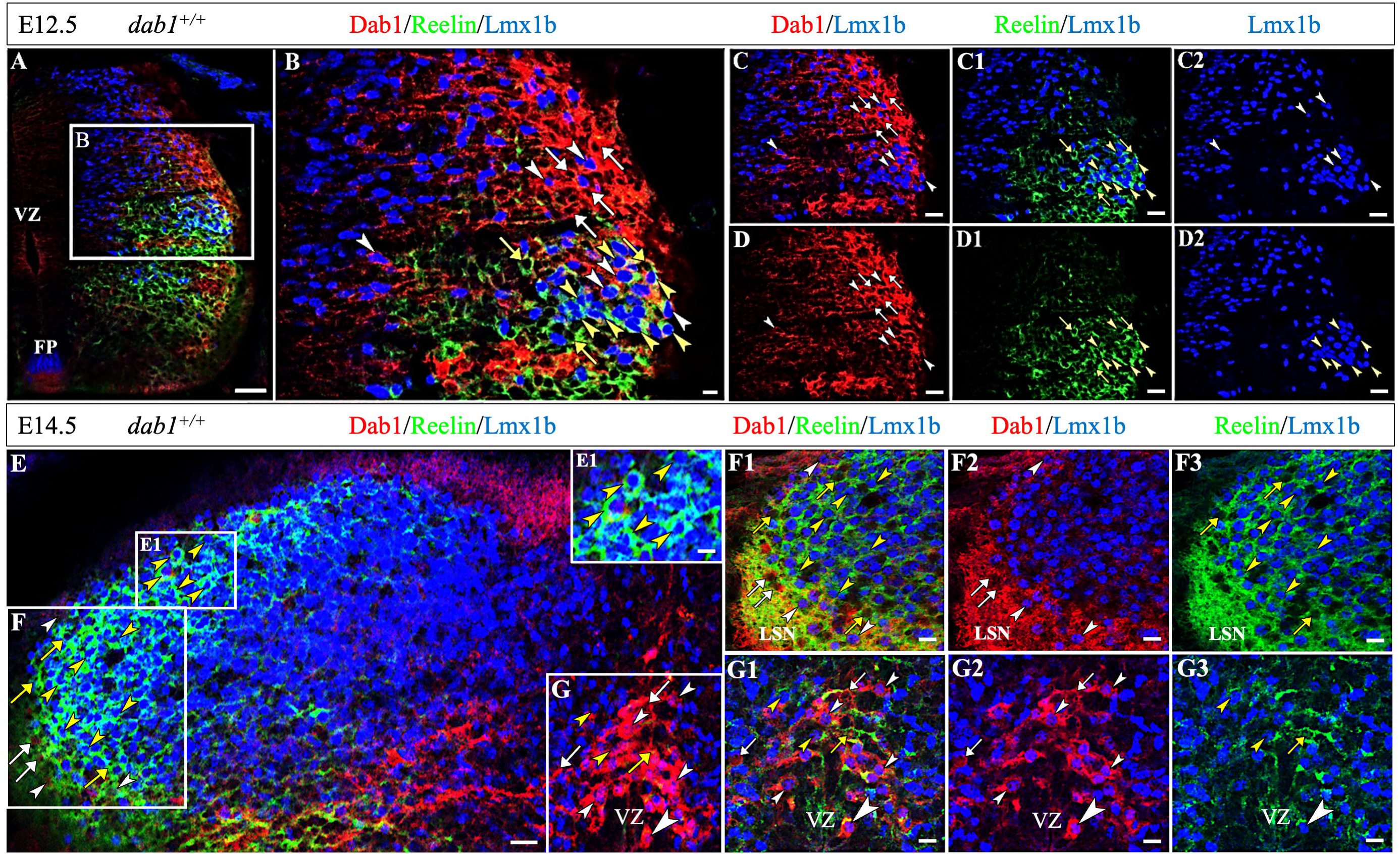
Dab1+Lmx1b+ and Reln+Lmx1b+ neurons appear as separate subsets of dI5 neurons. ***A***, An E12.5 hemisection shows localization of Dab1+Lmx1b+ (red with blue nuclei) and Reln+Lmx1b+ neurons (green with blue nuclei). Box in ***A*** is enlarged in ***B***, ***C-C2***, and ***D-D2***. ***B-D*,** Dab1+ (***B***, ***C***, ***D***, white arrows) and Dab1+Lmx1b+ neurons (***B***, ***C, C2, D,*** white arrowheads) are located dorsally compared to the more ventrally located Reln+ (***B***, ***D1***, yellow arrows) and Reln+Lmx1b+ neurons (***B, C1, D1***, ***D2,*** yellow arrowheads). Dab1+Lmx1b+ and Reln+Lmx1b+ neurons appear to be separate subsets of the Lmx1b+ dI5 neurons. ***E,*** An E14.5 section of a *dab1^+/+^*mouse triple-labeled with antibodies for Dab1, Reln, and Lmx1b. ***E1*** is enlarged in the upper right inset, and illustrates Reln+Lmx1b+ neurons (yellow arrowheads) within the superficial dorsal horn. Box ***F*** is enlarged in ***F1-F3*** and **G** is enlarged in ***G1-G3***. ***F-G,*** Both Dab1+ and Dab1+Lmx1b+ neurons are present in the lateral spinal nucleus (LSN) and the rim of the dorsal horn (***F1, F2,*** white arrows and arrowheads) together with Reln+ and Reln+Lmx1b+ neurons (***F1, F3,*** yellow arrows and arrowheads). Prominent Dab1+ (***G1, G2,*** white arrows) and Dab1+Lmx1b+ (***G1, G2***, white arrowheads) neurons are clustered just dorsal to the ventricular zone (VZ). Reln expression does not fill many of these neurons (Reln+ neurons, ***G1, G3,*** yellow arrow; Reln+Lmx1b+ neurons, ***G1, G3,*** yellow arrowheads). The Dab1+Lmx1b+ neuron at the large arrowhead in the ***G-G3*** appears to contain some Reln immunoreactivity. Scale bars: ***A,*** 50 µm; ***B,*** 20 µm; ***C-C2, D-D2,*** 30 µm; ***E,*** 40 µm; ***E1****, **F, F1-F3, G, G1-G3***, 25 µm.

At E14.5, Reln+Lmx1b+ and Dab1+Lmx1b+ neurons in the superficial dorsal horn do not appear to overlap, but it is difficult to interpret in the LSN due to the presence of secreted Reelin (Fig. 7F1-3). Dab1+ and Dab1+Lmx1b+ neurons are concentrated in a prominent midline group of neurons, just dorsal to the ventricular zone and interspersed with Reelin expression (Fig. 7G1-3). One of the E14.5 Dab1+Lmx1b+ neurons seemed to contain some Reelin within its cell body (Fig. 7G1, large arrowhead). Future studies are necessary to clarify if some of these neurons express both molecules or if the Reelin present in a few large Dab1+ neurons resulted from endocytosis (Hibi & Hattori, 2009; Morimura et al., 2005).

### Reln+Lmx1b+Zfhx3+ and Dab1+Lmx1b+Zfhx3+ dorsal spinal cord neurons are subsets of dI5 projection neurons

Based on single-cell RNA-sequencing, Osseward et al. (2021) showed that dI5/dIL_B_ neurons, like most cardinal classes, can be partitioned into group N and group Z neurons, with group-Z corresponding to long-range projection neurons marked by a transcription factor such as Zfhx3. We carried out triple immunolabeling experiments on wild-type mice to determine if Reln+Lmx1b+ neurons also express Zfhx3. A few Reln+Lmx1b+Zfhx3+ neurons were detected along the lateral rim of the *dab1^+/+^* superficial dorsal horn and in the LSN (Fig. 8A-C). At E13.5, Reln+Lmx1b+Zfhx3+ neurons were located laterally (Fig. 8A, red), and by E14.5 and E15.5 the Reln+Lmx1b+Zfhx3+ neurons were found more medially (Fig. 8B,C). The majority of Reelin+Lmx1b+ neurons were found in lamina II, an area composed predominantly of interneurons (Todd, 2010), and did not express Zfhx3 (Fig. 8).

**Figure 8:**
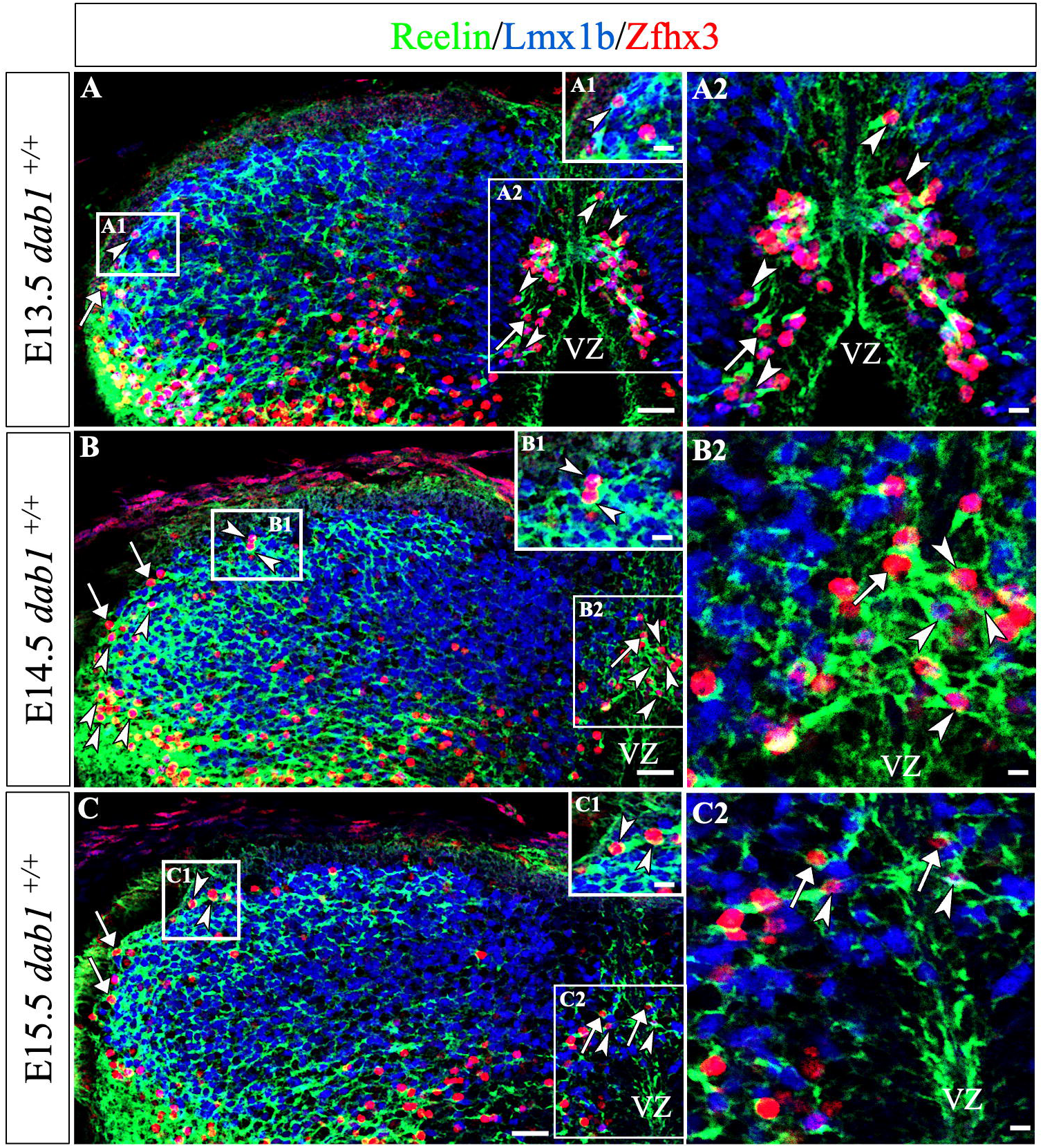
Large Reln+Lmx1b+Zfhx3+ neurons are projections neurons located along the rim of the dorsal horn and in the midline. Triple-labeled Reln+ (green), Lmx1b+ (blue nuclei) and Zfhx3+ (red nuclei) neurons (white arrowheads) from *dab1^+/+^* mice have green/cyan cytoplasm and purple nuclei (white arrowheads), compared to Reln+Zfhx3+ neurons with red nuclei (white arrows). ***A-A2,*** At E13.5, a triple-labeled neuron was detected laterally on the dorsal horn rim (***A1***, left box enlarged in inset) and others were found in the midline adjacent to the ventricular zone (VZ; ***A2*** box and enlargement, white arrowheads). ***B-B2,*** By E14.5, Reln+Lmx1b+Zfhx3+ and Reln+Zfhx3+ neurons are detected along the rim of the dorsal horn (***B1***, box enlarged in inset) and in the lateral spinal nucleus. Most neurons within the midline cluster are Reln+Lmx1b+Zfhx3+ neurons (white arrowheads), but a Reln+Zfhx3+ neuron (white arrow) is detected (***B2*** box and enlargement). ***C-C2,*** Examples of Reln+Lmx1b+Zfhx3+ and Reln+Zfhx3+ neurons are found along the E15.5 dorsal horn rim (***C1,*** box enlarged in the inset). While a few of the Reln+Lmx1b+Zfhx3+ neurons remain in the midline, most have migrated laterally (***C2*** box and enlargement). Scale bars: ***A-C,*** 40 µm; ***A1-C1,*** 25 µm; ***A2-C2***, 20 µm.

Data from E13.5 Dab1+Lmx1b+ dorsal spinal cords confirmed that some of these neurons also expressed Zfhx3 (Fig. 9A-C). The circumferential migration of lateral Dab1+Lmx1b+Zfhx3+ neurons progressed along the rim of the superficial dorsal horn between E14.5 (Fig. 9B) and E15.5 (Fig. 9C). Together, these results confirm that there are Reln+Lmx1b+Zfhx3+ and Dab1+Lmx1b+Zfhx3+ dI5 projection neurons along the rim of the superficial dorsal horn.

**Figure 9:**
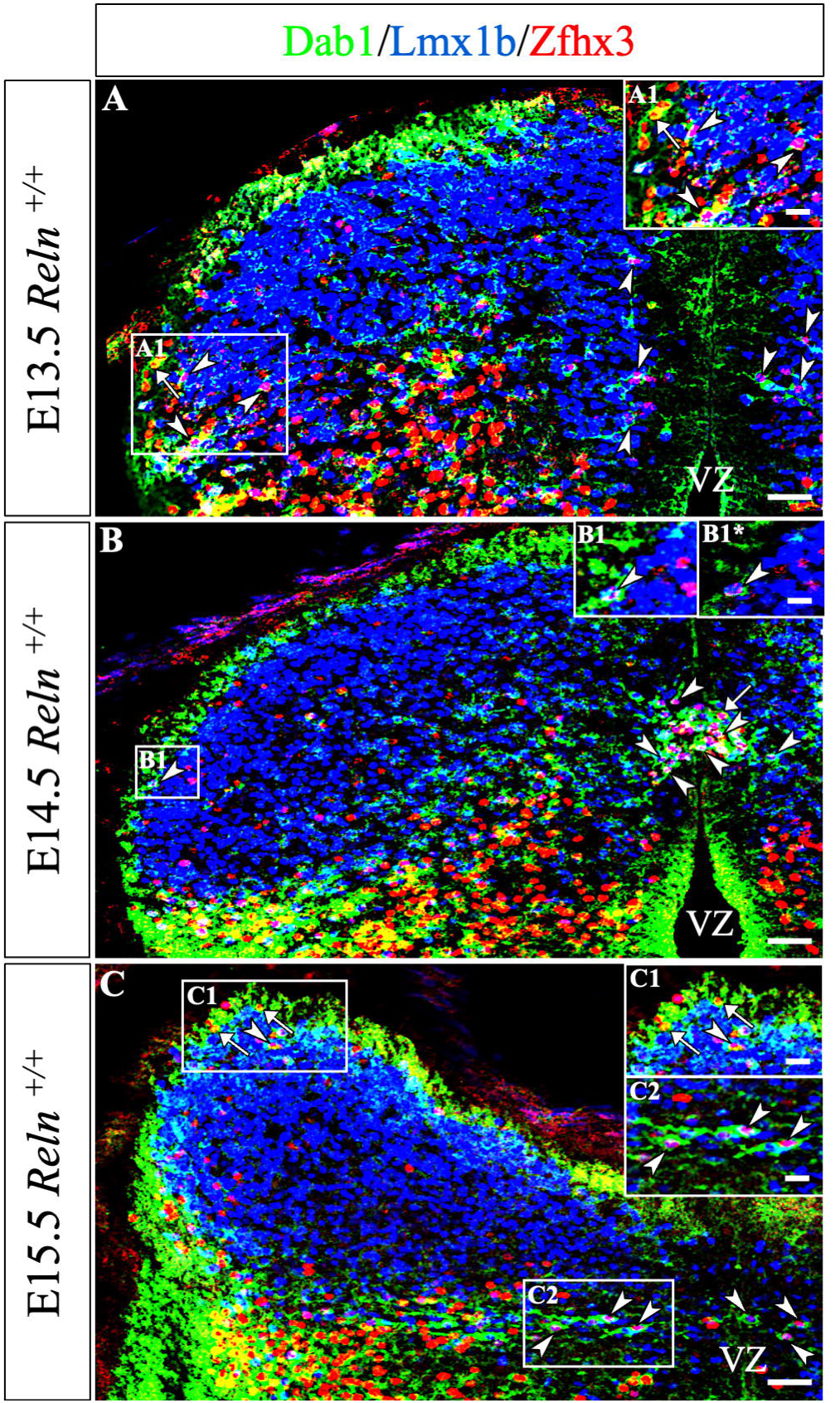
Dab1+Lmx1b+Zfhx3+ dI5 projection neurons migrate along the dorsal horn rim and from the midline into lamina V. Triple-labeled Dab1+ (green), Lmx1b+ (blue nuclei) and Zfhx3+ (red nuclei) neurons (white arrowheads) have green/cyan cytoplasm and purple nuclei. Examples of Dab1+Zfhx3+ neurons with red nuclei are marked by white arrows. ***A*,** Dorsal horn section has Dab1+Zfhx3+ neurons (white arrows, ***A1***, box enlarged in the inset) near the lateral border and in the lateral spinal nucleus. E13.5 Dab1+Lmx1b+Zfhx3+neurons (white arrowheads) are located near the midline above the ventricular zone (VZ). ***B,*** Example of a triple-labeled cell along the dorsal rim is enlarged from 2 different focal planes (***B1, B1*,*** white arrowheads, box enlarged in insets). By E14.5 the midline group above the VZ contains many Dab1+Lmx1b+Zfhx3+ neurons. ***C,*** By E15.5, more Dab1+Lmx1b+Zfhx3+ and Dab1+Zfhx3+ neurons are found along the dorsal horn rim (***C1,*** upper box enlargement,), whereas the medial dI5 neurons (***C2,*** lower box enlargement) have moved laterally into lamina V. Scale bars: ***A-C,*** 50 µm, ***A1, C1, C2,*** 30 µm; ***B1, B1*,*** 25 µm.

A number of Reln+Lmx1b+Zfhx3+ and Dab1+Lmx1b+Zfhx3+ neurons are associated with the dorsal midline and in lamina V (Figs. 8–9). Roome et al. (2020) identified a relatively early-born dI5 cluster of Paired-like homeobox 2A-expressing (Phox2a+) neurons that migrated toward the dorsal midline before they migrated laterally into lamina V. We found a number of triple-labeled Reln+Lmx1b+Zfhx3+ neurons near the midline at E13.5 (Fig. 8A, A2) and in the midline by E14.5 (Fig. 8B, B2). By E15.5, however, the number of Reln+Lmx1b+Zfhx3+ neurons in the midline fell (Fig. 8C2) and appeared to move further laterally into lamina V. A similar migratory pattern was observed for Dab1+Lmx1b+Zfhx3+ neurons. At E13.5 and 14.5, Dab1+Lmx1b+Zfhx3+ cells were located medially (Fig. 9A, B), and by E15.5, most had moved laterally (Fig. 9C). These results suggest that populations of Reln+Lmx1b+Zfhx3+ and Dab1+Lmx1b+Zfhx3+ neurons near the dorsal midline are dI5 projection neurons that migrate into lamina V.

To estimate the percentage of Zfhx3 neurons in the dorsal midline that express Reelin, we quantified them in E14.5 wild-type embryos. Of the total Zfhx3 neurons in the midline, we found that Reln+Lmx1b+Zfhx3+ neurons comprised 26 ±1%, Reln+ Zfhx3+ neurons were 5 ±3%, Lmx1b+Zfhx3+ neurons were 50 ±8%, and Zfhx3+ neurons represented 21 ±5% (rounded to whole cell). When we analyzed the percentage of midline Zfhx3 neurons that express Dab1, we found that Dab1+Lmx1b+Zfhx3+ cells make up 53 ±6% of total Zfhx3+ cells, while Dab1+Zfhx3+ compose 5 ±3%, Lmx1b+Zfhx3+ cells comprise 27 ±10%, and 14 ±1% of the Zfhx3+ neurons. Thus, Reln+Lmx1b+Zfhx3+ and Dab1+Lmx1b+Zfhx3+ are both subsets of dI5 projection neurons that migrate to and eventually settle in lamina V.

## Discussion

This study characterizes the migratory pathways of Reln+ and Dab1+ dorsal horn neurons relative to the glutamatergic Lmx1b-expressing early-born dI5 and late-born dIL_B_ neuronal subtypes. To our knowledge, this is the first example where Reln+ and Dab1+ neurons express the same neurotransmitter, are both mispositioned in the dorsal horn, and are influenced by the same transcription factors, Lmx1b and Zfhx3 (Cheng et al., 2004; Osseward et al., 2021; Szabo et al., 2015; Yvone et al., 2020; Yvone et al., 2017). The large, early-born Reln+Lmx1b+Zfhx3+ and Dab1+Lmx1b+Zfhx3+ dorsal horn neurons both migrate circumferentially along the rim of the dorsal horn, show evidence of subtle migratory errors, are subtypes of dI5 neurons, and express Zfhx3 which identifies them as projection neurons (Summary Fig. 10; Osseward et al., 2021; Roome et al., 2020; Sagner et al., 2021). A second, prominent group of dI5 Reln+Lmx1b+Zfhx3+ and Dab1+Lmx1b+Zfhx3+ neurons are found near the midline between E13.5 and E14.5, and then by E15.5, they are located laterally in lamina V. Based on their Zfhx3 expression, these triple-labeled cells are also long-range projection neurons (Osseward et al., 2021; Roome et al., 2020). In addition, at E14.5-15.5 there are Dab1+ and Dab1+Lmx1b+ neurons that migrate incorrectly across the deep dorsal horn of *Reln^-/-^* but not *Reln^+/+^* mice (Summary diagram, Fig. 10). Based on the lack of Zfhx3 labeling in lamina II of the superficial dorsal horn, the small Reln+Lmx1b+ and Dab1+Lmx1b+ neurons are likely interneurons and part of the dIL_B_ population. As dI5 neurons are integrated into adult pain circuits (Gross et al., 2002; Lai et al., 2016), the migratory errors sustained by these neurons likely contribute to the nociceptive abnormalities present in adult *Reln^-/-^*and *Dab1^-/-^*mice.

**Figure 10:**
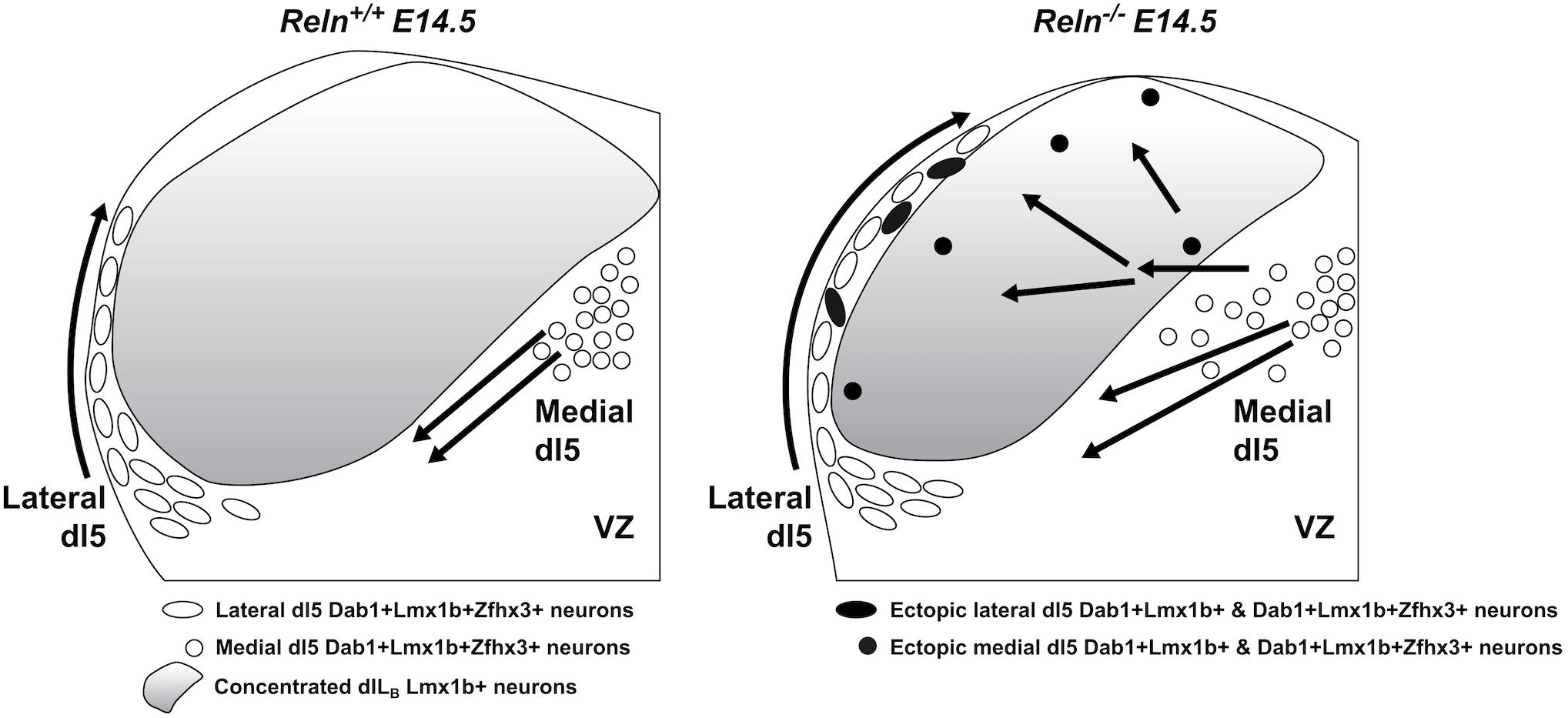
Summary Figure: Migratory errors of Dab1+Lmx1b+ and Dab+Lmx1b+Zfhx3+ neurons at E14.5. Early-born dI5 Dab1+Lmx1b+ and Dab1+Lmx1b+Zfhx3+ neurons display migratory errors in E14.5 *Reln^-/-^* compared to *Reln^+/+^*mice: 1) Lateral dI5 neurons migrate further dorsomedially along the *Reln^-/-^* dorsal horn rim and some are slightly mispositioned, 2) Medial dI5 neurons in *Reln^-/-^*dorsal horn form a disorganized midline cluster of Dab1+Lmx1b+Zfhx3+ cells. Some migrate tangentially across the deep dorsal horn, but most of these projection neurons settle in lamina V.

### Reln+Lmx1b+Zfhx3+ and Dab1+Lmx1b+Zfhx3+ dI5 neurons contribute to projection neurons in the superficial dorsal horn, LSN, and lateral lamina V

Based on birthdating studies, Altman & Bayer (1984) proposed that early-generated projection neurons (e.g., Waldeyer cells) migrate outwardly to the lateral edge of the embryonic spinal cord, turn dorsally, and then migrate circumferentially along the rim of the dorsal horn. Recent studies reported that early-born neurons (E9.5-10.5) that migrate along the dorsal rim, express the transcription factor Zfhx3, and were identified as projection neurons (Osseward et al., 2021; Sagner et al., 2021). In addition, Roome et al. (2020) reported a group of dI5 Lmx1b neurons that displayed a circumferential migration, and transiently expressed the Phox2a transcription factor in lamina I. These dI5 Phox2a commissural neurons were determined to be projection neurons that comprise part of the anterolateral system (Roome et al., 2020). Here we show that the Reln+Lmx1b+Zfhx3+ and Dab1+Lmx1b+Zfhx3+ neurons follow the circumferential pathway proposed by Altman & Bayer (1984) and that they are dI5 projection neurons.

Early-born dI5 neurons found in the dorsal midline co-expressed Zfhx3+ and EdU on E9.5-10.5 (Sagner et al., 2021) and migrated along the pattern reported in Roome et al. (2020) for the Lmx1b+Phox2a+ neurons into lamina V. These embryonic dI5-derived neurons migrate inward by E13.5, and then outward into lamina V by E15.5 (Roome et al., 2020), and our medial dI5 Reln+Lmx1b+Zfhx3+ and Dab1+Lmx1b+Zfhx3+ neurons follow a similar migratory pathway in wild-type mice. Phox2a^cre^ neurons are projection neurons (parabrachial and spinothalamic; Roome et al., 2020) based on their Zfhx3 expression. In addition, when Phox2a was conditionally deleted, Roome et al. (2020) observed the mispositioning of neurons in lamina V and the LSN, and migratory defects similar to those detected in Reln-signaling pathway mutants (Wang et al., 2012; Yvone et al., 2020; Yvone et al., 2017).

Alterations in the circumferential migratory pathway of the Dab1+Lmx1b+ dI5 neurons may underlie the positioning errors previously reported in the adult superficial dorsal horn and LSN. For example, adult *Reln^-/-^* and *dab1^-/-^* mice consistently have a 50% reduction of neurons in the LSN and lateral lamina V compared to wild-type controls (Akopians et al., 2008; Villeda et al., 2006; Wang et al., 2019; Yvone et al., 2020; Yvone et al., 2017). The presence of high Reln expression just lateral to the superficial dorsal horn may function to stop the migration of Dab1+ neurons when they reach their destinations. When part of the Reln-signaling pathway is absent, subsets of Dab1+ neurons fail to stop at their target destinations and instead continue along the circumferential pathway into the superficial dorsal horn.

Mispositioned Dab1+ sympathetic preganglionic neurons also failed to stop at their normal lateral location in thoracic spinal cord, and instead, migrated medially (Phelps et al., 2002; Yip et al., 2000). Krüger et al. (2010) suggested that when migrating sympathetic preganglionic neurons encountered Reln, Cofilin was phosphorylated, which in turn inhibited neuronal process extension and arrested migration. Thus, Reln expression in the spinal cord appears to stop the migration of Dab1+ neurons in the spinal cord when they reach their correct destinations.

### Migratory errors in *Reln^-/-^* E14.5 Dab1+ and Dab1+Lmx1b+ dorsal horn neurons

The first neuronal positioning errors identified in *Reln^-/-^* and *Vldlr^-/-^/ApoER2^-/-^*dorsal horns were large Dab1+ and Dab1+Neurokinin-1 Receptor-expressing (NK1R+) neurons found near the laminae II-III border that were not observed in *Reln^+/+^* and *Vldlr^+/+^/ApoER2^+/+^* mice (Akopians et al., 2008; Villeda et al., 2006). We also detected mispositioned Dab1+Lmx1b+NK1R+ neurons near the laminae II-III border of *Reln^-/-^* mice, and more Dab1+Lmx1b+NK1R+ neurons in *Reln^-/-^*laminae I-II than in *Reln^+/+^* mice (Wang et al., 2019; Yvone et al., 2020; Yvone et al., 2017). Furthermore, when mispositioned NK1R+ neurons, many of which are Dab1+, were selectively ablated in *dab1^-/-^* mice, these mutants lost their heat hypersensitivity but remained insensitive to mechanical stimuli (Wang et al., 2019). Here we show that some Dab1+ and Dab1+Lmx1b+ neurons in *Reln^-/-^*mice appear to dissociate from their radial glia prematurely, migrate tangentially across the E14.5 deep dorsal horn, and settle near the border of laminae II-III. A few of the dI5 neurons that migrate along the dorsal horn rim also appear incorrectly positioned just below lamina I. We suggest that these unusual migratory errors give rise to the mispositioned Dab1+ neurons near the laminae I-II and II-III borders in mutants and contribute to the thermal hypersensitivity of *Reln^-/-^* and *dab1^-/-^* mice (Akopians et al., 2008; Villeda et al., 2006).

Although the migratory errors of Dab1+ neurons in *Reln^-/-^* mice are more apparent than those of Reln+ neurons in *dab1^-/-^* mice, we detected mispositioned Reln+Lmx1b+ neurons in the adult superficial dorsal horn and the loss of Lmx1b+ neurons from the LSN and lateral reticulated area in the adult *dab1^-/-^* mice (Yvone et al., 2020). In addition, Szabo et al. (2015) demonstrated that the number of Reln-expressing cells in the superficial dorsal horn was significantly reduced following a conditional spinal cord deletion of Lmx1b, and that their mutants displayed mechanical insensitivity, similar to mutants of the Reln-signaling pathway. As both Reln+ and Dab1+ neurons are subsets of Lmx1b+ dI5 and are integrated into adult pain circuits, the migratory errors observed in Reln-signaling pathway mutants are the likely origins of the adult positioning errors that contribute to their nociceptive abnormalities.

## Acknowledgments

We thank Dr. Brian Howell for providing the *dab1^lacZ^* mice and Dab1 antibody, and Drs. Carmen Birchmeier and Thomas Müller for providing the Lmx1b antibody. We also thank several of our Reviewers for their important suggestions which improved our paper. This study acknowledges support from the National Science Foundation IOB-0924143 (PEP) and the Microscopy Core supported by NICHD of the National Institutes of Health under award number P50HD103557. We acknowledge support from the Eureka Scholarship, Hyde Fellowship, and Dissertation Year Fellowship to GMY, and support from NIH R25GM055052 (Hasson, PI) to CLCM.

## Data Availability Statement

The datasets and images generated and analyzed during the current study are available from the first and corresponding authors.

## Supplemental Figures

**Supplemental Fig. 1. Dab1+ axons in the medial marginal zone contribute to the dorsal columns (see Figure 4C, D).**

***A,*** The E14.5 *dab1^+/+^* dorsal horns contain Dab1 immunoreactivity (green, ***A-A1*)** in axons of the dorsal column (DC) pathway. The medial dI5 Dab1+Lmx1b+ neurons (green cytoplasm with red or purple nuclei, white asterisk in ***A***) are found just dorsal to the ventricular zone. In the Lmx1b (***A2***) and DAPI (***A3***) single channel images, the location of the axons is marked as DC. Scale bars: ***A-A3,*** 100 µm.

**Supplemental Fig. 2. Lateral Dab1+ and Dab1+Lmx1b+ neurons exhibit distinct positioning errors in E14.5 *Reln^-/-^* dorsal horn, enlarged from Figure 4C-D**.

Enlargements of E14.5 lateral dorsal horns of *Reln^+/+^* (***A-A2***) and *Reln^-/-^* mice (***B-B2***). ***A-B,*** Compared to *Reln^+/+^* mice, the Dab1+ (green, white arrows) and dI5 Dab1+Lmx1b+ neurons (green cytoplasm with red nuclei, lateral white arrowheads) in *Reln^-/-^* are positioned further medially along the rim of the dorsal horn (large white arrowheads) and there are disorganized cells along the superficial rim. In the *Reln^-/-^* deep dorsal horn (***B-B2***) distinct Dab1+ neurons (arrows) and Dab1+Lmx1b+ neurons (white arrowheads) migrate through the deep dorsal horn, whereas this area is almost devoid of Dab1+ neurons in *Reln^+/+^* mice (***A-A1***). Scale bars: ***A-A2, B-B2,*** 50 µm.

