## Supplementary figures and images for "Migration of dI5 Reelin-Lmx1b-Zfhx3 and Disabled-1-Lmx1b-Zfhx3 neurons contribute to the superficial dorsal horn and lamina V"

### Supplemental Figure 1

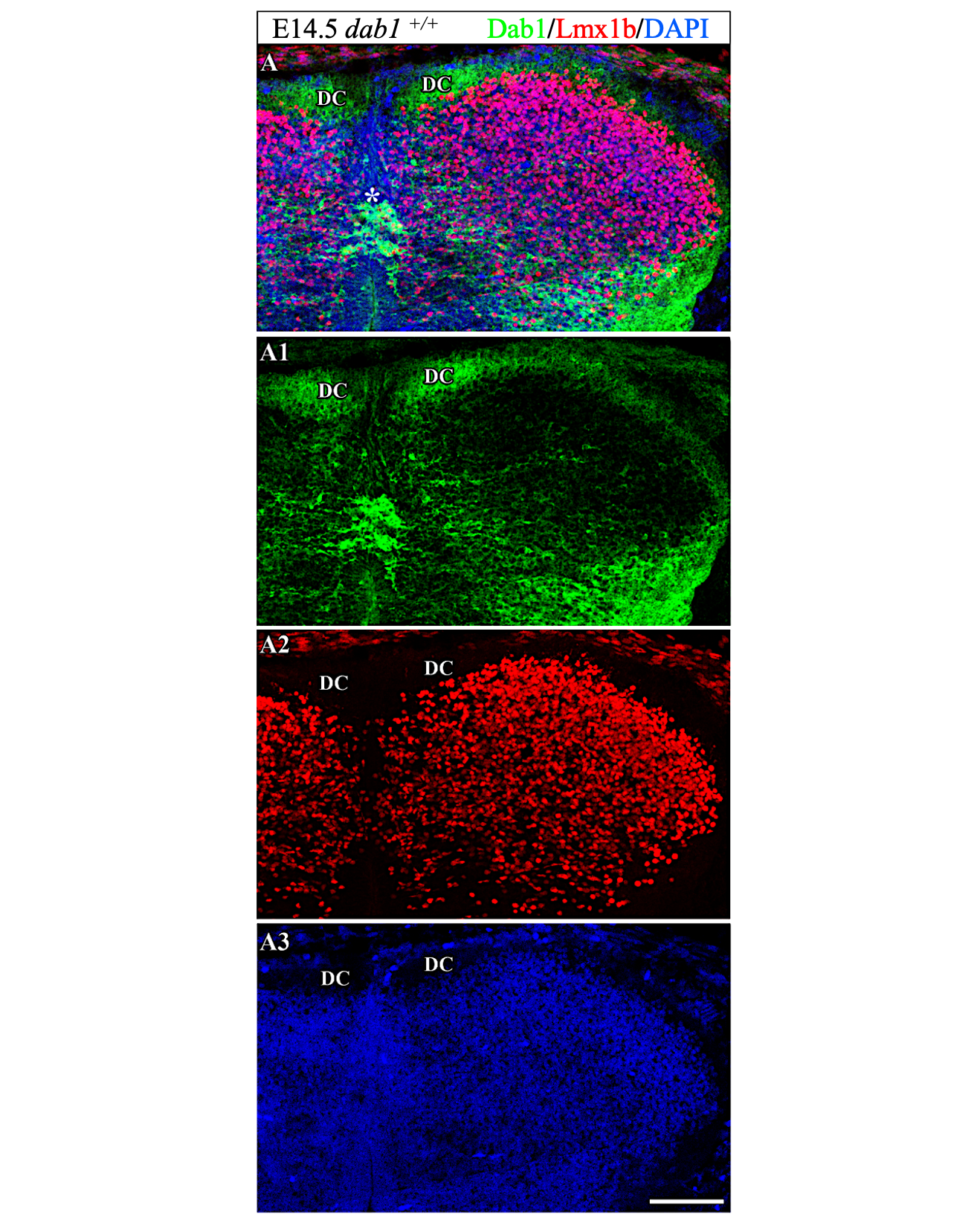

### Supplemental Figure 2

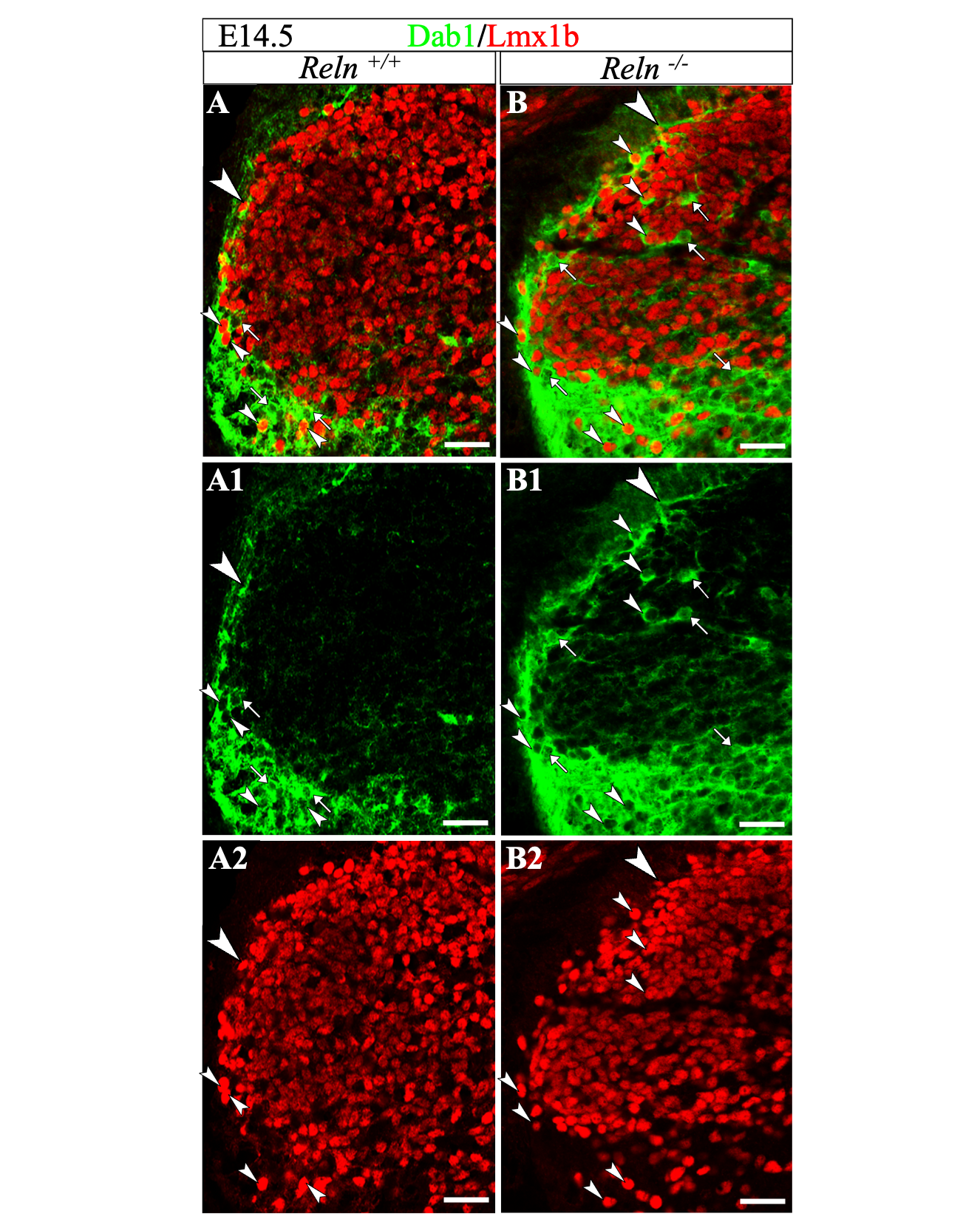
